# Benchmarking methods for genome annotation using Nanopore direct RNA in a non-model crop plant

**DOI:** 10.1101/2025.07.14.664836

**Authors:** Jade M Davis, Kristina K Gagalova, Lilian M V P Sanglard, Sabrina Cuellar, Mark R Gibberd, Fatima Naim

## Abstract

High-quality genome annotations are essential for transcriptomic analyses investigating plant responses to environmental stress. While Nanopore long-read direct RNA (dRNA) sequencing offers a powerful approach for improving genome annotations, studies benchmarking optimal tools for this process have primarily focused on animal models.

In this study, we benchmarked five annotation tools: StringTie2, IsoQuant, Bambu, FLAIR, and FLAMES, using dRNA data from barley infected with Net Form Net Blotch disease. We observed substantial variation across tools in isoform detection, structural completeness, splicing classification, and handling of 5′ read truncation. Several tools successfully identified novel transcripts, with the top-performing reference-guided approach detecting 994 previously unannotated transcripts, including candidates with predicted roles in disease response. Our results highlight the importance of plant-specific benchmarking of bioinformatic tools and demonstrate the utility of dRNA sequencing for improving genome annotations, supporting ongoing efforts to enhance reference resources for non-model plant species.

## Introduction

High-quality genome annotations that capture transcriptomic diversity are fundamental to many bioinformatic analyses, including differential gene expression and alternative splicing (AS) studies. While reference annotations are relatively well-curated for the model plant *Arabidopsis thaliana* [1], they remain limited for many non-model plant species [2,3]. Improving annotations is particularly challenging due to the inherent complexity of plant genomes, stemming from factors including polyploidy, large genome size, and high repeat content [4–7]. These factors necessitate substantial sequencing and computational effort to produce annotations that accurately define genic features, such as intron/exon boundaries and regulatory regions. Comprehensive annotations are especially critical for investigating AS, a key mechanism driving transcript and protein diversity in multicellular organisms, which has been increasingly linked to evolutionary adaptation [8]. AS modulates gene function and enhances phenotypic plasticity, shaping plant responses to both developmental cues and environmental stressors [9]. It has been implicated in responses to abiotic stressors such as drought, salinity, and low temperature [10–15], as well as to biotic challenges including viral and fungal infections [16–19].

Several computational strategies can be employed to improve plant genome annotations [20], with transcriptome-based approaches playing a central role in structural annotations. By capturing actively expressed transcripts, RNA sequencing (RNA-seq) enables annotation refinement that reflects transcriptomic variation across specific conditions or tissue types. For example, in cassava (*Manihot esculenta*), annotation enrichment using RNA-seq under cold stress facilitated the identification of 323 additional differentially expressed genes compared to using the reference Ensembl annotation [21]. Other studies have enhanced genome annotations for various plants including *Brassica oleracea* (wild cabbage), *Zea mays* (maize), *Papaver somniferum* (opium poppy), and *Amaranthus hypochondriacus* (prince’s-feather) [22–25].

Third-generation long-read RNA-seq technologies, such as those developed by Oxford Nanopore Technologies (ONT) and Pacific Biosciences (PacBio), offer significant advantages for RNA-based annotations [26]. These technologies can generate reads spanning thousands of bases [27], enabling full-length transcript capture. In contrast, conventional short-read platforms, which typically produce reads ≤300 bp [28], often lack the resolution necessary to reconstruct complete transcript isoforms [29]. Among long-read methods, ONT’s direct RNA (dRNA) sequencing, which selectively captures native polyadenylated mRNA without PCR amplification, has proven particularly effective for detecting full-length and low-abundance transcripts. This approach has uncovered novel isoforms in rice, bamboo, and *Arabidopsis thaliana* [30–32], including approximately 38,500 previously unannotated transcripts in the latter, proving a powerful tool for enriching plant genome annotations.

A growing number of bioinformatic tools support dRNA-directed annotation, including StringTie2, IsoQuant, Bambu, FLAIR, and FLAMES. Each employs distinct algorithms for transcriptome assembly, from splice or intron graphs to read clustering and correction workflows (Table 1) [33–37]. The diversity of approaches creates uncertainty about which tools are best suited to specific datasets or annotation goals. Accordingly, several recent benchmarking efforts have evaluated long-read annotation tools, notably, the Long-read RNA-Seq Genome Annotation Assessment Project (LRGASP) [38]. The study compared both reference-guided and *de novo* annotation strategies across real and simulated datasets from human, mouse, and manatee. Other studies have also benchmarked tools in humans and model organisms such as mouse, *Drosophila*, and *Caenorhabditis elegans* [39,40]. Tool performance varied across sequencing technologies, reference quality and overall annotation strategies, highlighting a potential need for plant-specific benchmarking [41].

**Table 1.** Comparison of long-read transcriptome annotation tools.

| Tool | Approach | Key Features | Reference Annotation |
| --- | --- | --- | --- |
| **StringTie2**<br>[33] | Splice graph | Builds exon-splice graphs; selects paths with the highest read support | Optional |
| **IsoQuant**<br>[34] | Intron graph | Simplifies intron-based graphs; finds best-supported transcript paths | Optional |
| **Bambu** [35] | Read clustering + novel discovery rate (NDR) | Clusters reads; uses NDR to filter novel transcripts, pretrained model fallback | Optional |
| **FLAIR** [36] | Align–Correct–Collapse | Corrects junctions with short-read or reference data; iterative filtering | Recommended for correction |
| **FLAMES**<br>[37] | Read clustering + collapsing | Collapses similar/truncated isoforms; optimised for single-cell, but supports bulk | Required |

Barley, a globally important cereal crop, has a large and highly repetitive genome [42], making it an ideal system for benchmarking dRNA annotation tools. While recent efforts have vastly improved barley annotations [43–47], such as PanBaRT20 which captures 79,600 genes and 582,000 transcripts across multiple tissues and cultivars [48], existing resources mainly reflect developmental and abiotic stress conditions. Host responses to biotic stressors, including Net Form Net Blotch (NFNB) disease of barley, caused by *Pyrenophora teres* f. *teres*, remain underrepresented. Although NFNB-induced gene expression has been studied [49–52], standard barley genome reference annotations were used. AS under disease pressure is also poorly characterised, primarily due to inadequate annotations. Accordingly, long-read dRNA sequencing could be applied to enrich annotations, although plant-specific benchmarking of annotation tools is still needed. This study addresses the gap by benchmarking reference-guided and *de novo* dRNA annotation tools in barley and assessing the biological relevance of novel transcripts identified in NFNB disease of barley.

## Systems and methods

### Plant and fungal propagation and inoculation of leaves

Methodology for fungal inoculation and leaf tissue harvest was adapted from Moolhuijzen et al. (2023) as follows [53]. Barley (cv. RGT Planet) was grown for 3 weeks under natural light in glasshouse conditions (Curtin University). A fungal spore solution was prepared using frozen plugs from *Pyrenophora teres* f. *teres* (NB29 isolate [54]). A 10 μL spore solution droplet (2000 spores/mL) was placed 10 cm and 15 cm from the tip of the second mature leaf. The detailed protocol for spot inoculation is available online [55].

### Sample collection, RNA extraction and sequencing

RNA extraction was performed as described in Moolhuijzen et al. (2023). Three biological replicates for control and NFNB-infected leaf samples were harvested 3 days post-inoculation (dpi). Infected leaves were harvested using a 2 mm biopsy punch, with three punches collected at observed disease lesions (above, within and below lesion) in a line parallel to the midrib. Control samples were harvested by targeting a similar leaf area. Each replicate consisted of three punches taken from two lesions per leaf from four individual leaves, resulting in a total of 24 punches. Total RNA was extracted using the Plant/Fungi Total RNA Purification Kit (Norgen Biotek, Canada) and eluted in 40 μL. Samples were quantified using a Nanodrop spectrophotometer (NanoDrop Technologies, USA) and a Qubit™ RNA High Sensitivity (HS) kit run on a QuantiFluor RNA system E3310 (Qubit, UK).

RNA samples were sequenced using both Illumina short-read RNA sequencing and Oxford Nanopore Technologies (ONT) long-read direct RNA (dRNA) sequencing. Illumina library preparation and sequencing were performed at Genomics WA using the SureSelect XT HS2 mRNA Library Preparation Kit (Agilent, USA), followed by 2 × 150 bp paired-end sequencing on the NovaSeq 6000 platform (Illumina, USA). This generated approximately 32-37 million raw read pairs per replicate. For dRNA sequencing, equal amounts of RNA from biological replicates for each treatment were pooled, and 1000 ng of RNA was used for library preparation with the ONT Direct RNA Sequencing Kit (SQK-RNA004, ONT, UK). To increase transcript diversity in the control barley dRNA library, three additional RNA samples were included; these were generated using the same preparation protocol. The only modification to the manufacturer’s protocol was reducing the RNA Calibration Strand (RCS) solution volume to 0.1 μL. Libraries were loaded onto two FLO-PRO004RA flow cells and sequenced using the ONT PromethION 2 Solo platform.

### Direct RNA read pre-processing and mapping

Direct RNA reads were used to generate annotations for tool comparison. Unless otherwise specified, all bioinformatic tools were run with default parameters. Raw pod5 files were basecalled using Dorado v0.7.3 [56] with the rna004_130bps_sup@v5.0.0 model. RCS sequences were removed by aligning reads to the ENO2 reference (Ensembl transcript ID YHR174W_mRNA, accessed August 2024 [57]) using Minimap2 v2.28-r120955 [58] with recommended ONT dRNA settings (-ax splice -uf -k14). Unmapped reads were extracted with Samtools fastq v1.21 (-f 4) [59] and filtered using Chopper v0.9.0 [60] with minimum quality and length thresholds set at Phred 10 and 100 bp (-q 10 -l 100). Quality control was performed using FastQC v0.12.1 [61] and MultiQC v1.25.2 [62]. Cleaned reads were aligned to the RGT Planet reference genome [46] using Minimap2 v2.28-r120955 with recommended ONT dRNA settings. Pre-processing yielded 3.6 million reads (99.9% alignment) for the control sample and 6.1 million reads (96.1% alignment) for the NFNB-infected sample. Alignments were sorted and indexed using samtools v1.21 [59].

### Transcript discovery and annotation

Five tools were evaluated for reference-guided annotation: StringTie2, FLAIR, IsoQuant, Bambu and FLAMES [33–37]. All analyses used the RGT Planet annotation [46] in the respective input parameter. StringTie2 v3.0.0 was run with the long-read (-L) parameter and the output for both samples was combined with the reference annotation using --merge. For FLAIR v2.0.0, BAM files were converted to BED format using FLAIR bam2Bed12 and refined using FLAIR correct with the reference annotation. Refined BED files were concatenated before processing with FLAIR collapse, applying the -- annotation_reliant flag. Bambu v3.8.3 was run with quant = FALSE to omit quantification. IsoQuant v3.6.3 was run with --data_type nanopore, and the SAMPLE_ID.extended_annotation.gtf output was used. The 925331f GitHub commit of FLAMES (https://github.com/mritchielab/FLAMES; accessed February 2025) was run using the ‘bulk_long_pipeline’ workflow. Configuration followed example documentation (https://mritchielab.github.io/FLAMES/reference/bulk_long_pipeline.html, accessed February 2025), except barcode demultiplexing and quantification were omitted and no_flank was set to false. For *de novo* annotation, Bambu, StringTie2, and IsoQuant were executed as above without reference-guided parameters. Bambu was tested with novel discovery rate (NDR) thresholds of 0.1, 0.5, 0.75, and 1. NDRs between 0.1 and 0.75 produced errors to increase NDR, meaning a value of 1 was selected.

Annotations were standardised using AGAT agat_convert_sp_gxf2gxf.pl v1.4.1 [63] and cleaned with GenomeTools GFF3 v1.6.5 [64] with -sort and -tidy options. Coding sequences (CDS) were annotated with GenomeTools CDS v1.6.5 with -startcodon -matchdescstart flags, and untranslated regions (UTRs) were managed using AGAT agat_sp_manage_UTRs.pl v1.4.1.

### Annotation quality analysis and benchmarking

Gene and transcript counts were obtained using AGAT agat_sp_statistics.pl v1.4.1. Coding potential was assessed by extracting transcripts with gffread v0.12.7 [65] and classifying them using RNAsamba classify v0.2.5 [66] with the full_length_weights.hdf5 model. Transcript splicing was classified using a custom Python script (available at 10.5281/zenodo.15636556). Transcript-level precision and sensitivity were calculated against RGT Planet using GffCompare v0.12.6 [65] with -R to skip reference annotations not overlapped by query models. Novel splice junctions and transcript structures were analysed with SQANTI3 v5.3.6 sqanti3_qc.py [67] using --short_reads with raw Illumina reads. Following Dong et al. (2023) [68], junctions with more than ten uniquely mapping reads were considered validated. Visualisations were generated in RStudio 2024.12.0+467 [69] using ggplot2 v3.5.1 [70] and RColorBrewer v1.1-3 palettes [71].

### Visualisation of transcript models

To assess annotation quality in *de novo* approaches, ten transcripts associated with disease resistance expressed in the NFNB-infected sample were visualised. To select these transcripts “disease resistance” was queried in the functional annotation search of the panBARLEX portal (https://panbarlex.ipk-gatersleben.de/, accessed February 2025) and results for RGT Planet were extracted. Expression was assessed using IsoQuant v3.6.3 with --no_model_construction --data_type nanopore parameters and the RGT Planet annotation. Ten transcripts with more than 20 mapped reads were randomly selected for visualisation.

To assess annotation quality in reference-guided approaches, five of the disease resistance-associated transcripts and four novel transcripts were visualised. Novel transcripts were identified by querying annotations against RGT Planet using GffCompare v0.12.6. For each tool, a random transcript labelled “unknown” (intergenic to all reference annotations) was selected, except for FLAMES, which reported none. Selected annotations from both *de novo* and reference-guided approaches were visualised alongside RNA alignments using IGV v2.17.3 [72].

### Functional characterisation, visualisation and expression analysis of novel transcripts

Novel transcripts identified by StringTie2 (reference-guided) were functionally characterised. Transcripts labelled “unknown” by GffCompare v0.12.6 against RGT Planet were extracted using AGAT agat_sp_extract_sequences.pl v1.4.1 run with –merge and queried against the PanBaRT20 [48] protein database (https://ics.hutton.ac.uk/panbart20/downloads/PanBaRT20_transuite_transfeat_pep_renamed.fasta.gz, accessed March 2025) using BLASTx v2.12.0 [73]. Transcripts with >95% percentage identity and query coverage were classified as known. A custom script was used to extract annotations for novel transcripts, followed by conversion into a protein FASTA using AGAT agat_sp_extract_sequences.pl v1.4.1 with -t cds -p --cfs –cis parameters. Functional domains were predicted with InterProScan v5.73-104.0 with the v5.73-104.0 data release using --disable-precalc -goterms. Gene Ontology (GO) terms were extracted, counted and then summarised using REVIGO v1.8.1 [74] (http://revigo.irb.hr/, accessed March 2025) with the “higher value is better” setting, a small list and *Triticum aestivum* set as the closest reference. The biological processes treemap was exported and then refined using ggplot2 v3.5.1 and treemap v2.4-4 in RStudio 2024.12.0+467. Novel transcripts with predicted functions were visualised with read alignments in IGV v2.17.3 [72] to assess transcript structure. Transcripts were classified as true if supported by ≥3 splice-aligned reads and false if not. Transcripts substantially shorter than alignments were also classified as false (see Supplementary Figure 1).

Expression of novel transcripts was assessed using short-read Illumina data. Raw reads were processed with nf-core/rnaseq v3.18.0 [75] using STAR/RSEM, CSI indexing and the StringTie2 reference-guided annotation (--aligner star_rsem --bam_csi_index). Transcript counts from rsem.merged.transcript_counts.tsv were analysed in DESeq2 v1.46.0 [76] using a Wald test. Transcripts with log2 fold change ≥2 and adjusted p-value ≤ 0.005 were considered significantly differentially expressed. Finally, dplyr v1.1.4 [77] was used to identify novel significantly differentially expressed transcripts.

### Data and code availability

Raw read sequencing data is available on the National Centre for Biotechnology Information Sequence Read Archive under the BioProject ID PRJNA1282305. Code for all analysis, comparison and visualisations performed can be accessed at 10.5281/zenodo.15636533. These commands include embedded use of a Nextflow-based [78] pipeline to streamline processing, available at <u>10.5281/zenodo.15636556</u>.

## Implementation and results

### Annotation tools performed variably

Annotation output varied substantially across tools. Among reference-guided methods, StringTie2 annotated the most genes and transcripts (53,929 genes/58,119 transcripts), followed closely by IsoQuant (53,444/57,825) and Bambu (52,081/54,695), all of which exceeded the RGT Planet reference annotations. In contrast, counts were considerably lower for FLAIR (28,073/28,979) and FLAMES (11,694/20,041). For *de novo* approaches, Bambu annotated the highest number of genes (16,463), followed closely by StringTie2 (16,027) and lastly IsoQuant (13,661). Bambu reported a substantially higher transcript count (50,587) than IsoQuant (15,105) and StringTie2 (18,046) (Figure 1A).

**Figure 1.**
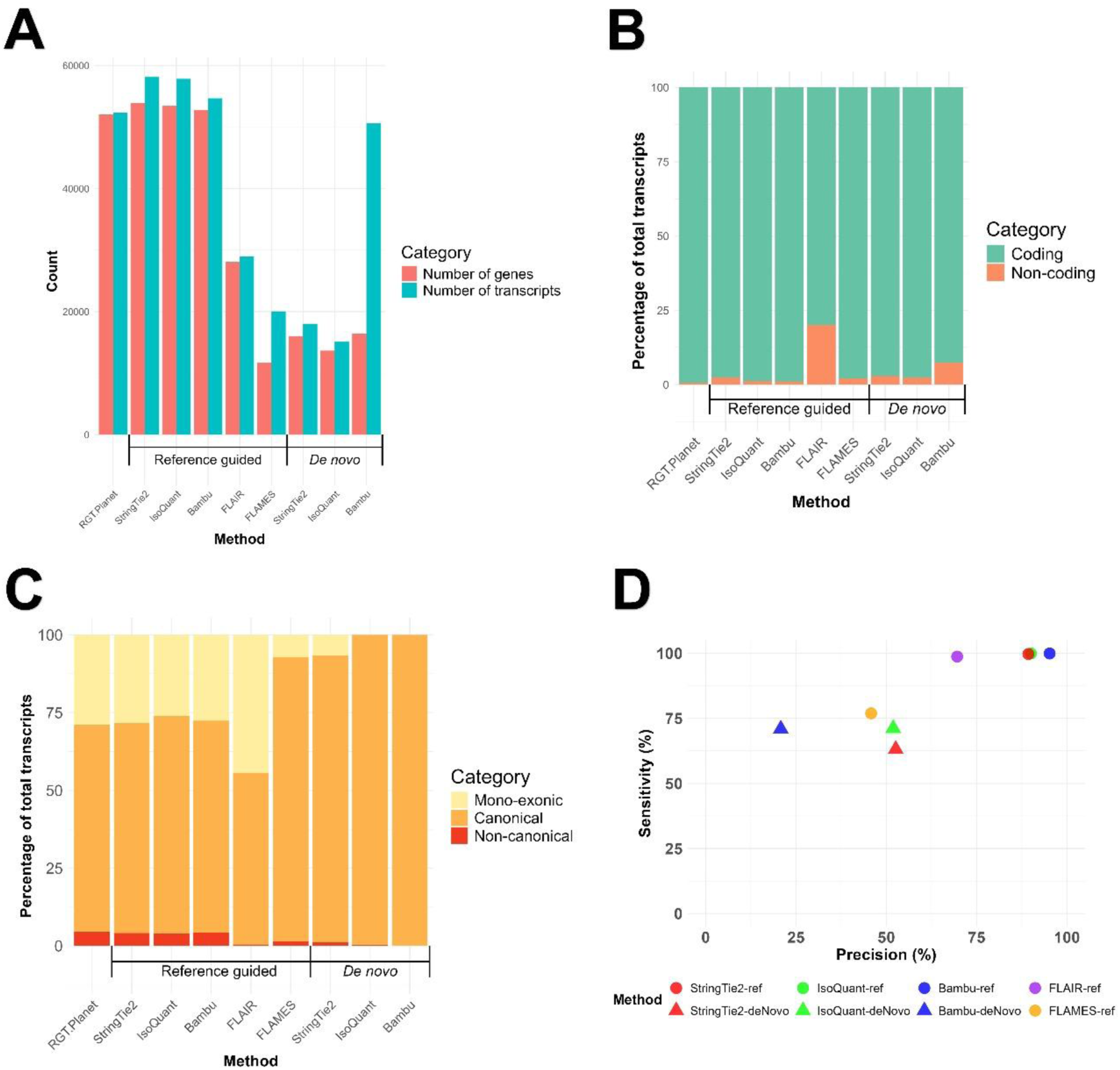
Comparative metrics to benchmark reference-guided and *de novo* methods for genome annotation of barley (cv. RGT Planet) against reference annotations of RGT Planet. **(A)** Summary of transcript and gene counts generated for annotation tools **(B)** analysis of transcript annotations for coding potential**, (C)** splicing patterns, and **(D)** transcript-level precision and sensitivity values compared to reference annotations of RGT Planet.

Predicted coding potential, for reference-guided methods, was consistent with the reference for StringTie2, IsoQuant, Bambu, and FLAMES (<3% non-coding), while 20% of FLAIR transcripts were non-coding. In *de novo* mode, IsoQuant and StringTie2 maintained <3% non-coding transcripts, while Bambu reached 7% (Figure 1B).

Splicing profiles differed between tools, especially for mono-exonic transcripts, which are rare in eukaryotic genomes and often indicative of pseudogenes [2]. Reference-guided annotations from StringTie2, IsoQuant and Bambu showed splicing profiles similar to reference annotations. FLAMES produced a lower percentage of mono-exonic transcripts and a higher percentage of canonically spliced transcripts, while FLAIR annotated a higher proportion of mono-exonic transcripts. Among *de novo* methods, IsoQuant and Bambu produced no mono-exonic transcripts, whereas StringTie2 contained 6% (Figure 1C).

Transcript-level precision and sensitivity also varied. In reference-guided methods, Bambu, StringTie2, IsoQuant and FLAIR achieved high sensitivity (>98%), while FLAMES ranked lowest (77%). More variation was observed for precision, with Bambu ranking highest (95%), followed by IsoQuant (90%) and StringTie2 (89%), with FLAIR (70%) and FLAMES (46%) much lower. In *de novo* methods, IsoQuant and Bambu had similar sensitivity (∼71%), with StringTie2 slightly lower (63%). Precision was highest for IsoQuant and StringTie2 (∼52%), while Bambu was lowest (21%) (Figure 1D).

### Transcript structure and validation

SQANTI3 classification revealed insights into isoform diversity. Full splice matches (FSMs, known annotations) made up the majority of annotations across all tools and methods, except Bambu *de novo*, which was dominated (53%) by incomplete splice matches (ISMs, transcripts with matching splice junctions to known transcripts but missing exons). Notably, 98% of these Bambu *de novo* ISMs were 3′ fragments, meaning 5′ exons were missing. Most annotation sets, except for FLAMES, included intergenic transcripts (transcripts from novel genes). All except FLAIR included a notable number of novel not in catalogue (NNC) transcripts, denoting annotations with at least one novel splice junction within a known gene. Other structural categories, such as genic (overlapping known exons/introns), and novel in catalogue (NIC, novel combinations of known splice sites) represented a smaller proportion (Figure 2A). Intron retention, the most common alternative splicing event in plants [79], was highest in FLAMES (415 transcripts), followed by IsoQuant (339), StringTie2 (170), FLAIR (68) and Bambu (9), in reference-guided annotations.

**Figure 2.**
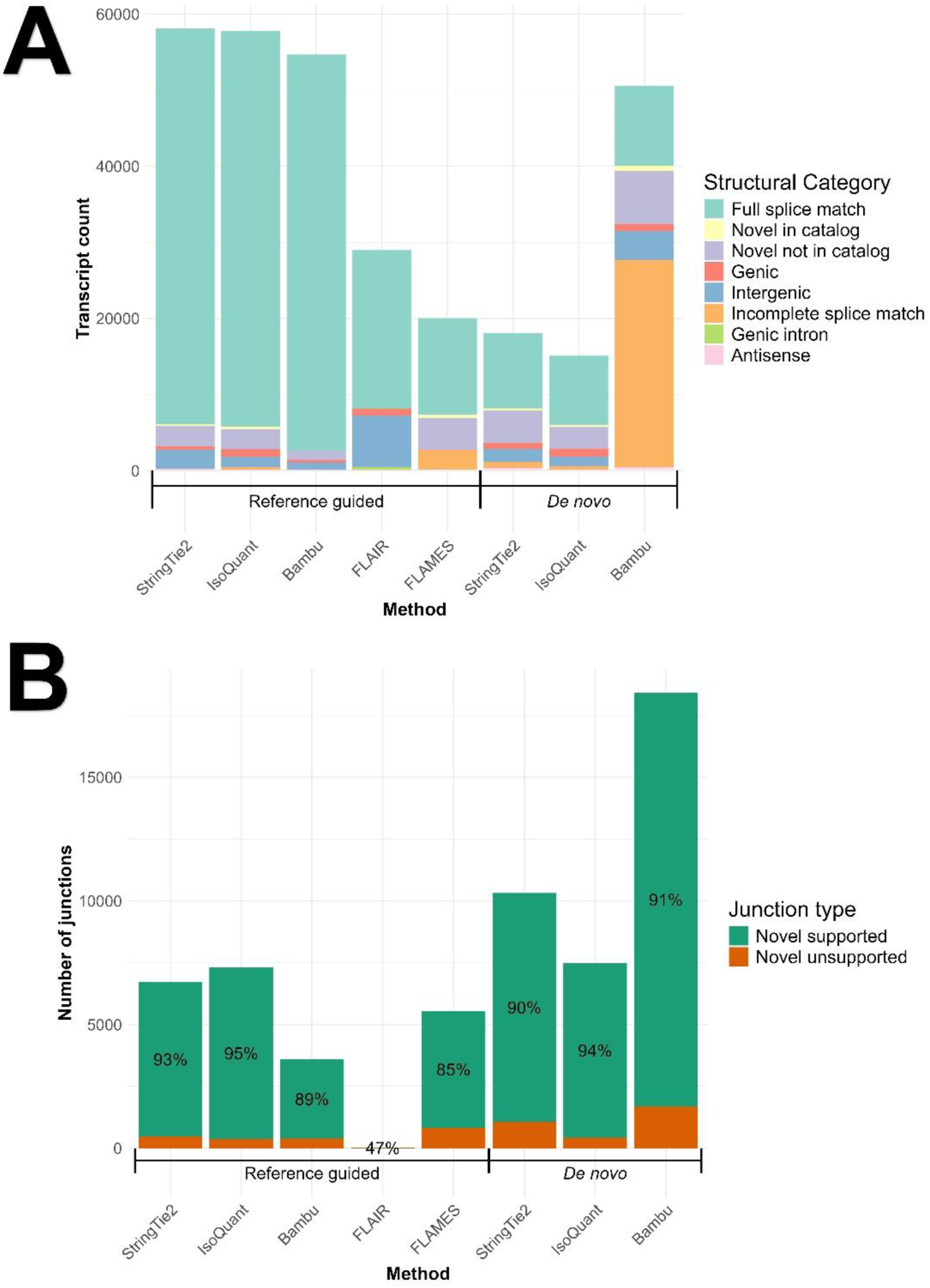
SQANTI3 analysis of annotations generated for barley (cv. RGT Planet) with reference-guided and de novo methods. **(A)** Structural category of transcripts compared to the RGT Planet reference annotation, and **(B)** short-read support for novel splice junctions, the percentage of novel junctions supported is shown on the bars.

Validation of novel junctions with Illumina short reads showed support across most methods and tools. IsoQuant and StringTie2 had the highest support among reference-guided methods (95% and 93%, respectively), followed by Bambu (89%) and FLAMES (85%). Despite over 20% of FLAIR transcripts being intergenic, only 15 novel junctions were identified (47% Illumina support). All novel FLAIR junctions were within known genes, showing that all intergenic annotations were mono-exonic. Novel junctions annotated by *de novo* methods showed high Illumina support, led by IsoQuant (94%) and closely followed by Bambu (91%) and StringTie2 (90%) (Figure 2B).

### Visualisation reveals tool-specific annotation patterns

Annotations and read alignments were visualised for qualitative evaluation of transcript models, revealing several tool-specific patterns. Some tools, particularly Bambu *de novo*, annotated transcripts with identical 3′ intron/exon structure to known transcripts which differed in the length/count of 5′ exons, typically with low 5′ read coverage (Figure 3A). IsoQuant sometimes annotated additional genes and transcripts at loci with existing annotations (Figure 3B).

**Figure 3.**
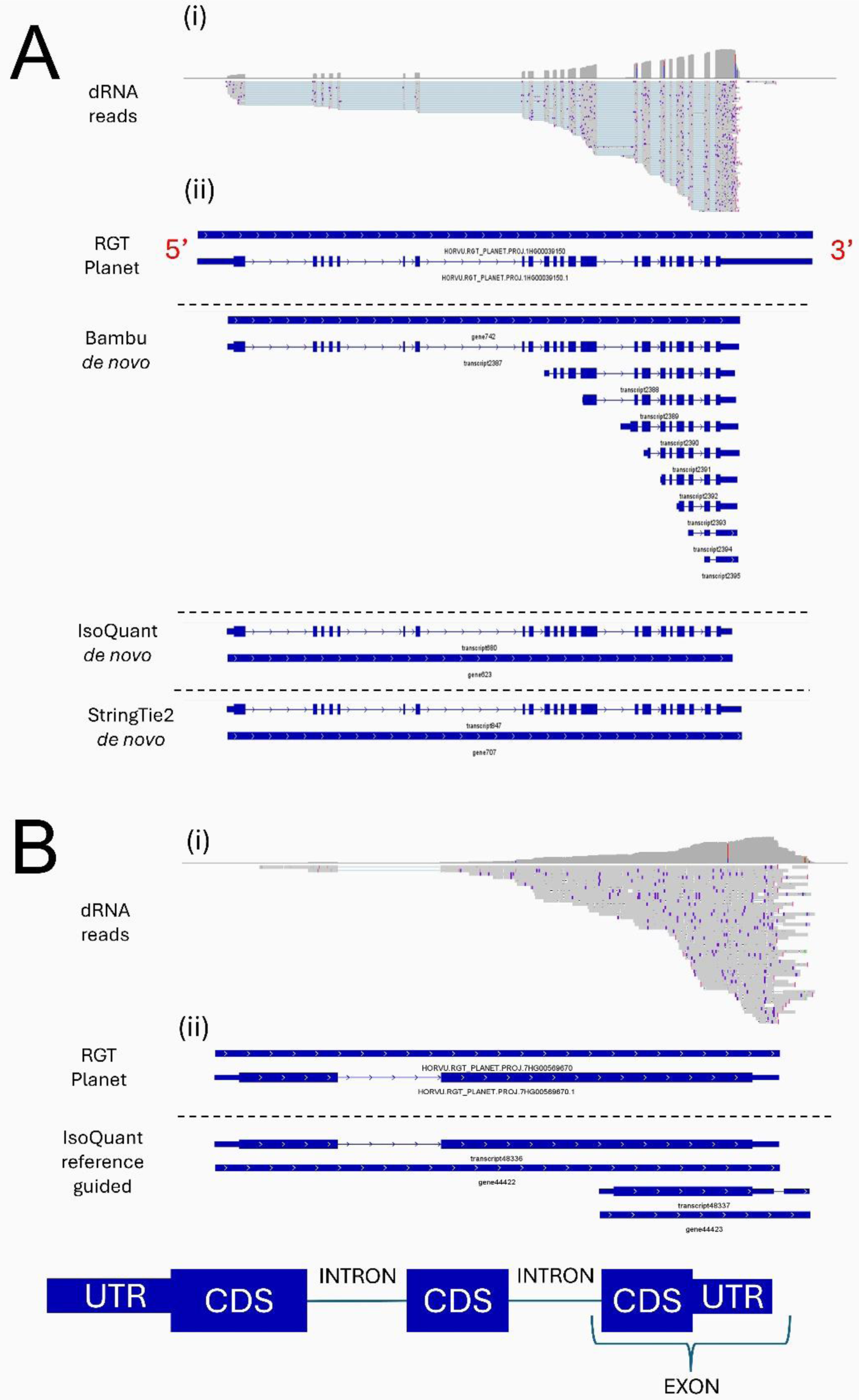
Representative examples of two patterns seen in visual investigation of annotations produced using reference guided and *de novo* methods. **(A)** Abundance of annotations with identical splicing at the 3′ end, differing in the length of their 5′ exon for Bambu *de novo* and not supported by reference annotations or StringTie2 and IsoQuant *de novo*, and **(B)** Duplicated transcript models at loci with existing annotations generated by IsoQuant. (i) Read alignments with corresponding coverage profiles, (ii) representative reference transcript models. In gene annotations, thinner lines represent introns and thicker lines represent exons, split into coding sequences (CDS) and untranslated regions (UTR). Arrows point from the 5′ to 3′ direction.

Reference-guided annotations were visualised at four novel and five disease resistance-associated genes. StringTie2 and IsoQuant captured all novel genes, though both missed 5′ exons at one locus. IsoQuant also introduced an extra gene and transcript at one locus. Bambu annotated three novel genes (one missing a 5′ exon), FLAIR annotated two (both missing 5′ exons), and FLAMES did not annotate any novel genes (Supplementary Figure 2A). For resistance-associated genes, every tool except FLAMES generated models at all five loci. StringTie2 and Bambu matched the reference without issues, while FLAIR and IsoQuant annotated additional 3′ fragments and genes, respectively. FLAMES annotated transcripts at three loci, all with extra 5′ and/or 3′ truncated transcripts.

For *de novo* annotations, ten resistance-associated genes expressed in the diseased sample were visualised. StringTie2 successfully annotated full-length transcripts at eight loci, while IsoQuant annotated transcripts at seven, including one with extra gene annotations and another lacking complete resolution. Bambu also annotated eight loci but generated additional 5′ truncated transcripts at four loci and failed to fully resolve two transcripts (Supplementary Figure 3).

### Novel transcript discovery and functional characterisation

Based on strong performance, the reference-guided StringTie2 annotation was selected for further analysis. Comparison with RGT Planet and PanBaRT20 identified 994 previously unannotated transcripts. Of these, 74 had at least one Gene Ontology (GO) term predicted by InterProScan. REVIGO summarisation revealed involvement in several biologically important functions, including stress response, receptor signalling, proteolysis, and transcription regulation (Figure 4). Manual inspection showed that 74% of novel functionally annotated transcripts were supported by ≥3 reads; 9% were classified as false as their models were missing several exons at the 5′ end of read alignments. Short-read RNA-seq analysis identified 6,791 significantly differentially expressed (DE) transcripts, 258 of which were previously unannotated.

**Figure 4.**
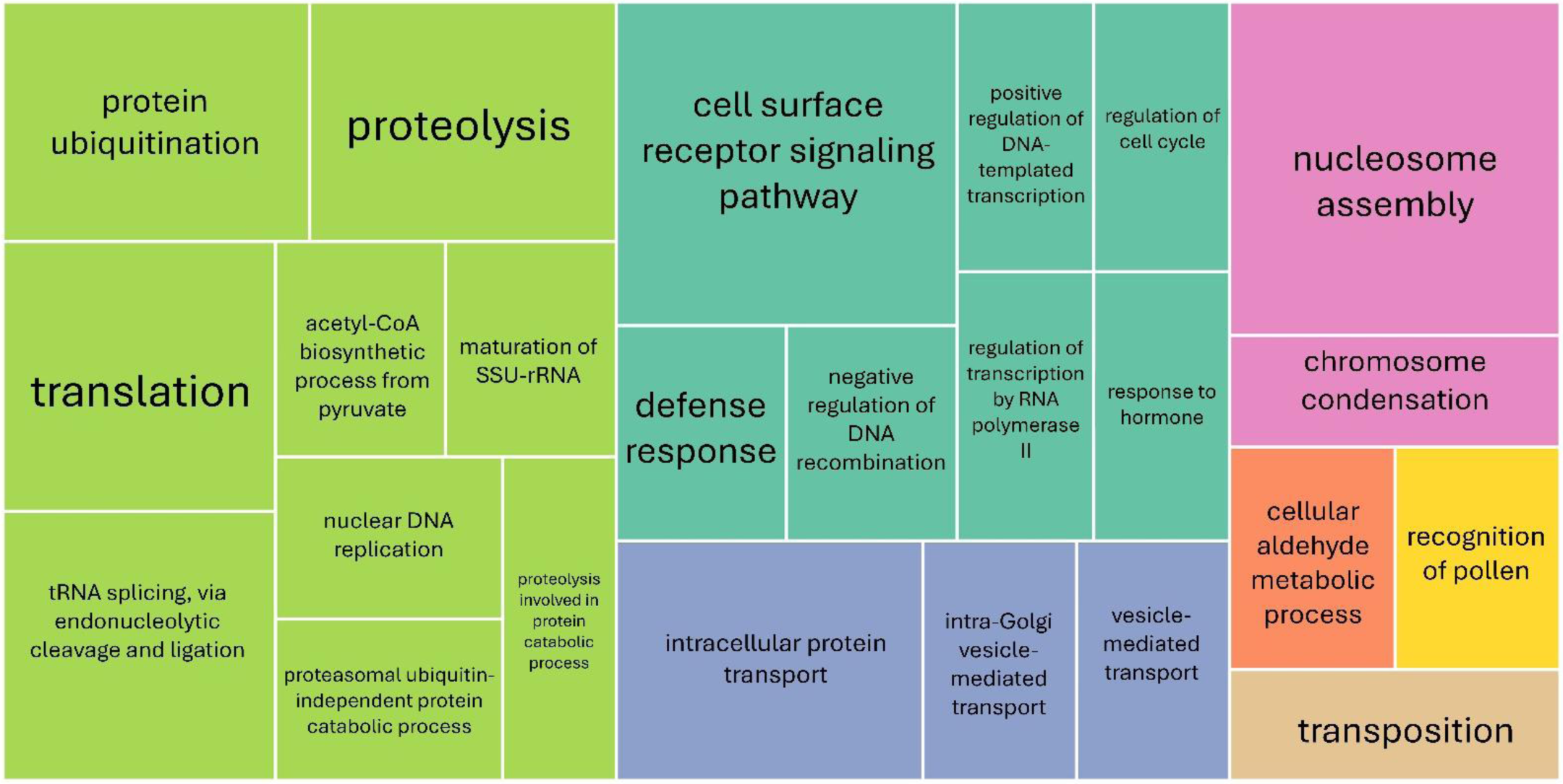
Tree map showing predicted gene ontology (GO) terms for novel transcripts identified by StringTie2 using barley direct RNA data. Functional annotation analysis for novel transcripts performed by InterProScan and REVIGO was used to generate a tree map of GO terms. Box sizes correspond to the GO term frequency, and colours correspond to terms classed by REVIGO as ‘similar’.

## Discussion

This study benchmarked five bioinformatic tools for barley genome annotation using direct RNA (dRNA) sequencing, applying both reference-guided and *de novo* approaches. While previous benchmarking efforts have focused on model organisms, this work provides a crop plant-specific perspective. Comparisons to LRGASP [38] and two other benchmarking studies [39,40], offered valuable reference points for evaluating tool performance.

### 5′ truncation is a major limitation of dRNA data

A consistent challenge across tools was apparent 5′ transcript truncation. Bambu *de novo* frequently annotated transcripts with identical 3′ ends to known transcripts with missing 5′ exons, supported by SQANTI3’s detection of a high proportion of ISMs. Visualisation of StringTie2 reference-guided annotations also showed that 9% lacked 5′ exons despite read support. This reflects a known limitation of ONT dRNA sequencing, which initiates sequencing of the RNA-DNA hybrid molecule from the 3′ poly-A tail, leading to stronger 3′ coverage and tapering at the 5′ end. As read coverage declined towards 5′ ends, annotation tools likely misclassified truncated reads as independent isoforms. The challenge of ONT 5′ truncation has been discussed in previous studies [80], and was further noted in LRGASP, which reported that tools generally demonstrated greater transcript 3′ resolution. The limitation was also noted by Su et al. (2024), who simulated ONT datasets with complete reads and datasets with 5′ and/or 3′ read truncation. The study reported overall decreases in annotation precision and sensitivity for most tools using truncated datasets, supporting the negative impact of incomplete reads on annotation quality.

Several strategies could be trialled in future work to mitigate the impact of 5′ truncation. Firstly, the use of alternative reverse transcriptase (RTs), may optimise the process to ensure RNA-cDNA hybrids are representative of full-length transcripts. Additionally, an increase in flow cell output may improve full-length transcript detection. Given the immense size of the barley transcriptome [44], deeper sequencing is likely needed to achieve sufficient transcript coverage, which may be facilitated by continued improvement of ONT technology into the future. The use of 5′ adapters could also be explored to improve transcript resolution. This technology replaces 5′ caps with an identifiable oligomer, allowing for full-length read detection; it has been previously applied for full-length transcript capture in plant, viral and human transcriptomes [81–83]. More stringent filtering for minimum read length during pre-processing may also be beneficial, although this introduces the risk of excluding true short transcripts. Addressing 5′ bias remains a key priority for improving dRNA-informed annotations.

### StringTie2 is a top reference-guided performer

Analysis of reference-guided approaches revealed significant differences between tools. Overall, FLAMES and FLAIR performed less reliably. FLAIR, for example, annotated a large proportion of intergenic, mono-exonic transcripts, a finding supported by a 2025 study, which identified similar issues using FLAIR with ONT cDNA data from human brain samples [84]. FLAIR also exhibited poor reliability in other analyses, annotating few genes and transcripts, with a high percentage of non-coding predictions. Sagniez et al. (2024) also concluded that FLAIR was not as successful as other tools. FLAMES, on the other hand, did not annotate any intergenic transcripts or novel genes, a result consistent with Su et al. (2024), who tested FLAMES on 16 long-read datasets and found no intergenic transcripts. Furthermore, FLAMES was less sensitive and precise, with a higher percentage of ISM models. Interestingly, Su et al. (2024) noted that the quality of reference annotations had a greater impact on the performance of FLAIR and FLAMES than other tools. Our results further support this finding and demonstrate the unsuitability of FLAIR and FLAMES for the annotation of complex crop genomes, such as barley, which lack reference resources for comprehensive conditions.

In contrast, StringTie2, IsoQuant, and Bambu performed relatively well in reference-guided mode, showing high precision and near-perfect sensitivity. High sensitivity was expected for IsoQuant and Bambu, which automatically output reference annotations regardless of expression. While StringTie2 was run to merge the reference annotation with the assembled transcripts, achieving similar high sensitivity. All three tools produced annotations with coding and splicing profiles similar to RGT Planet, and thus, these metrics were not a viable basis for comparison. IsoQuant revealed limitations in handling read truncation and Bambu, while strong overall, fell behind StringTie2 in gene annotation, splice junction support, and performance in visualisations. These findings highlight StringTie2 as a top performer for reference-guided dRNA annotation in non-model plants.

### StringTie2 and IsoQuant are top *de novo* performers

*De novo* annotation comparisons revealed greater variation. Bambu annotated a significant proportion of 3′ fragment ISMs, likely due to the high novel discovery rate (NDR = 1), which retained all read classes, leading to the discovery of ‘novel’ transcripts informed by truncated reads. Therefore, Bambu’s model for *de novo* annotation, trained on human data [35], appears to be unsuitable for barley. However, Bambu does allow users to train models on species-specific data, which could improve its performance, although this was beyond the study’s scope. StringTie2 and IsoQuant performed more consistently, with similar precision, sensitivity, and SQANTI3 profiles. IsoQuant omitted mono-exonic transcripts due to default ONT settings that suppress unspliced transcripts to avoid false positives; this setting can be changed by users if desired. IsoQuant slightly outperformed StringTie2 for Illumina junction support, while StringTie2 performed better in visual comparisons, showing strengths for each tool. Therefore, both are viable options for *de novo* annotation in barley.

### StringTie2 enables discovery of novel, disease-related transcripts

The use of StringTie2 enabled the discovery of 994 novel transcripts, 74 of which were functionally annotated. Predicted roles included several biologically important functions, notably stress response, receptor signalling, proteolysis and transcriptional regulation [85–87], demonstrating the value of dRNA-based annotation enrichment under disease conditions. However, 9% of novel transcripts lacked significant annotation of 5′ exons, reinforcing that even top-performing tools are still constrained by dRNA’s technical limitations. Although StringTie2 successfully enriched the annotation of disease-relevant transcripts, correcting 5′ truncation will be essential for maximising annotation quality.

## Conclusion

While this study provides novel insights into dRNA-informed barley annotation, several limitations should be noted. Firstly, this study focused only on default or recommended annotation parameters and used minimal read filtering (Phred ≥10; length ≥100 bp). However, more stringent thresholds or tool-specific tuning may improve results. Furthermore, unlike other benchmarking efforts, this study did not include spike-in controls, which may assist in evaluating the quality of dRNA data and tool performance for isoform detection [88]. Importantly, tool selection alone does not guarantee comprehensive annotations, and informed curation is required to ensure that plant transcript models reflect biological truth and not just computational artefacts [89]. Best practices include integrating both short- and long-read data, using multiple tools, filtering mono-exonic transcripts lacking protein domains and more routine visualisation to determine the accuracy of annotations [2]. These strategies will help reduce artefacts and enhance annotation reliability.

This study offers novel insights into dRNA-based genome annotation in barley and demonstrates the value of StringTie2 for discovering biologically meaningful transcripts under disease pressure. Our findings could inform improvements to barley reference resources and guide dRNA annotation strategies in other non-model crops. Future efforts could be supported by using the Nextflow-based pipeline provided with the analysis code, which streamlines dRNA pre-processing, annotation and analysis. Continued advances in long-read sequencing and tool development will be essential for building complete, functional annotations to support crop improvement and agrigenomic research.

## Author contributions

FN and MRG conceptualised the study. FN designed the laboratory experiment. JMD, LN and SC performed the laboratory experiment. JMD and KKG conceptualised and conducted the benchmarking bioinformatics experiment, including code and pipeline development and performed data analysis. JMD wrote the first draft of the manuscript with FN and KKG. All authors contributed to the review and final manuscript.

## Acknowledgments

The computational component of this research was conducted using the high-performance computing resources of the Pawsey Supercomputing Research Centre’s Setonix Supercomputer (https://doi.org/10.48569/18sb-8s43). We thank Ashley Jones, Simon Dunbar, Alka Saxena, and Johannes Debler for valuable discussions about the use of Oxford Nanopore Technologies and RNA sequencing.

