## Supplementary figures for "Benchmarking methods for genome annotation using Nanopore direct RNA in a non-model crop plant"

### Supplementary material

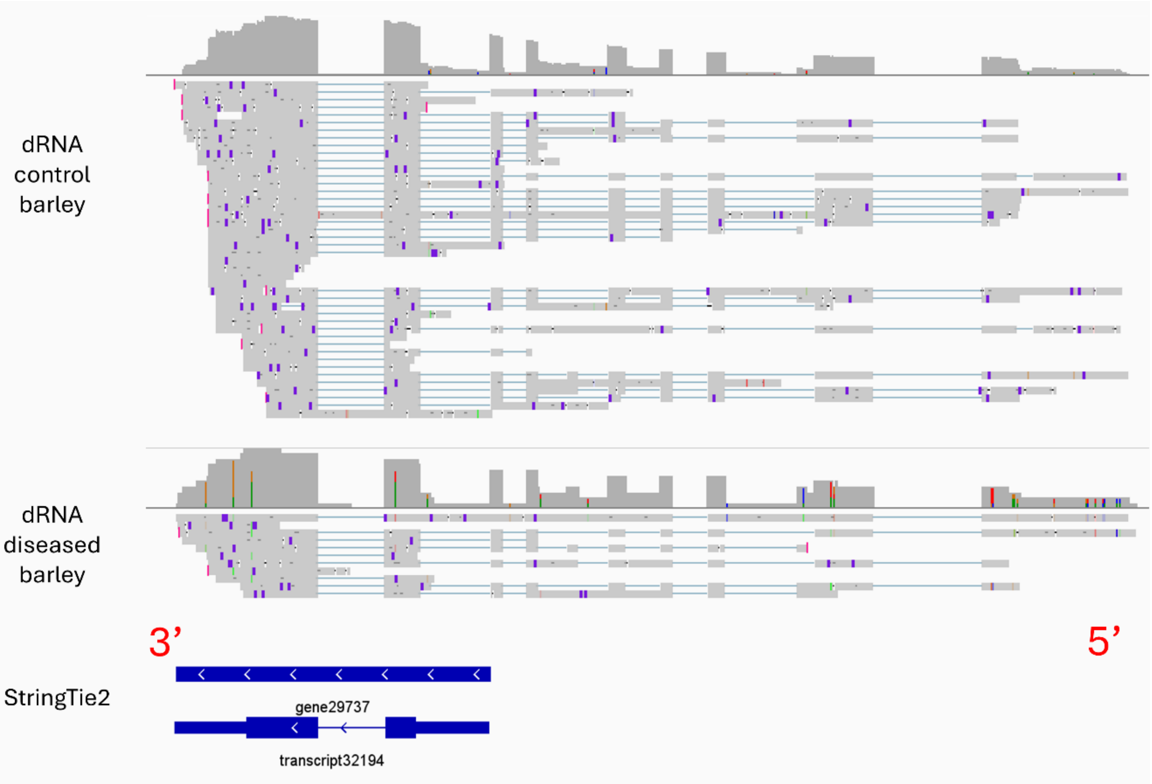


**Supplementary Figure 1.** Representative example of novel transcripts identified by StringTie2 reference-guided mode, which did not cover a large proportion of 5' alignment lengths.

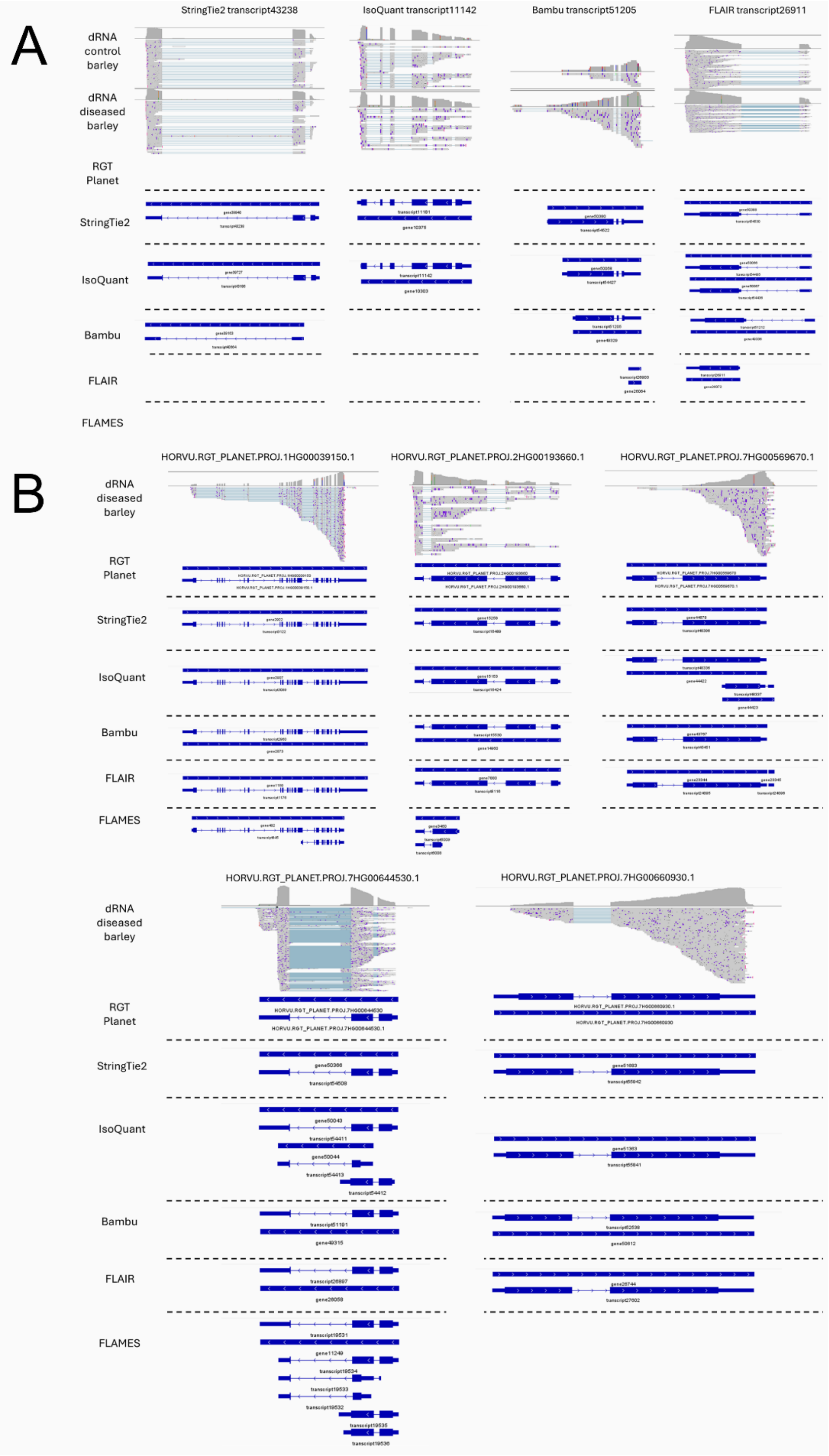


**Supplementary Figure 2.** IGV visualisation of annotations produced using annotation tools guided by the RGT Planet reference annotation and direct RNA (dRNA) reads from uninfected and net-form net-blotch (NFNB) infected barley; **(A)** four genes novel to RGT Planet **(B)** five resistance-associated genes.

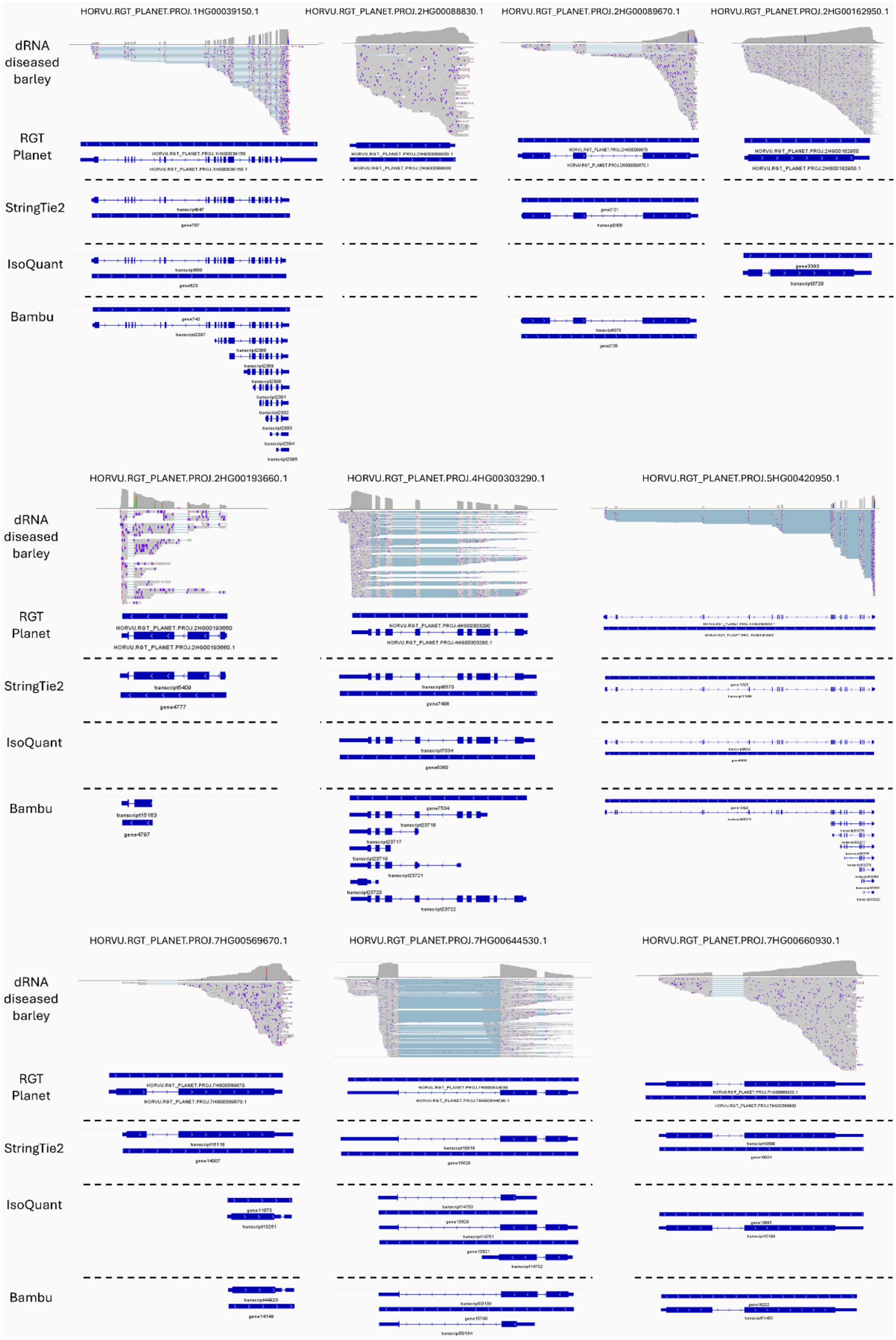

**Supplementary Figure 3.** IGV visualisation of *de novo* annotations produced using different programs with direct RNA (dRNA) reads from uninfected and net-form net-blotch infected (NFNB) barley as input.
